# Activated PI3Kδ syndrome, an immunodeficiency disorder, leads to sensorimotor deficits recapitulated in a murine model

**DOI:** 10.1101/2021.01.15.426862

**Authors:** Ines Serra, Olivia R. Manusama, Fabian M. P. Kaiser, Izi Izumi Floriano, Lucas Wahl, Christian van der Zalm, Hanna IJspeert, P. Martin van Hagen, Nico J.M. van Beveren, Sandra M. Arend, Klaus Okkenhaug, Johan J.M. Pel, Virgil A.S.H. Dalm, Aleksandra Badura

**Affiliations:** Department of Neuroscience, Erasmus MC, Rotterdam, The Netherlands; Department of Immunology, Erasmus MC, Rotterdam, The Netherland; Department of Pediatrics, Erasmus MC, Rotterdam, The Netherlands; Division of Clinical Immunology, Department of Internal Medicine, Erasmus MC, Rotterdam, The Netherlands; Department of Psychiatry, Erasmus MC, Rotterdam, The Netherlands; Department of Infectious Diseases, Leiden University Medical Center, Leiden, The Netherlands; Department of Pathology, University of Cambridge, Cambridge, United Kingdom; Academic Center for Rare Immunological Diseases (RIDC), Erasmus MC, Rotterdam, The Netherlands

**Author notes:** **Correspondence**: Aleksandra Badura, Department of Neuroscience, Erasmus MC, Wytemaweg 80, 3015 CN Rotterdam. **Conflict of interest statement:** PMH has received grants and research support from Takeda, CSL Behring, Abbvie, Lamepro, Novartis Nederland, and honoraria or consultation fees from UCB Pharma. The other authors have no conflict of interest to declare.

## Abstract

The phosphoinositide-3-kinase (PI3K) family plays a major role in cell signalling and is predominant in leukocytes. Gain-of-function (GOF) mutations in the *PIK3CD* gene lead to the development of activated PI3Kδ syndrome (APDS), a rare primary immunodeficiency disorder. A subset of APDS patients also displays neurodevelopmental delay symptoms, suggesting a potential role of *PIK3CD* in cognitive and behavioural function. However, the extent and nature of the neurodevelopmental deficits has not been previously quantified. Here, we assessed the cognitive functions of two APDS patients, and investigated the causal role of the *PIK3CD* GOF mutation in neurological deficits using a murine model of this disease. We used E1020K knock-in mice, harbouring the most common APDS mutation in patients. We found that APDS patients present with visuomotor deficits, exacerbated by autism spectrum disorder comorbidity, whereas p110δ^E1020K^ mice exhibited impairments in motor behaviour, learning and repetitive behaviour patterning. Our data indicate that *PIK3CD* GOF mutations increase the risk for neurodevelopmental deficits, supporting previous findings on the interplay between the nervous and the immune system. Further, our results validate the knock-in mouse model, and offer an objective assessment tool for patients that could be incorporated in diagnosis and in the evaluation of treatments.

## Introduction

Primary immunodeficiencies (PID) encompass a group of heterogeneous, mostly inheritable, disorders that affect distinct components of the immune system (1,2). Common manifestations of PID include increased susceptibility to infection, autoimmune disease, auto-inflammatory complications and malignancies, ultimately leading to increased morbidity and mortality rates (3–6). Activated PI3K delta (PI3Kδ) syndrome (APDS) is a rare monogenic PID, caused by heterozygous mutations in either the *PIK3CD* or *PIK3R1* genes, encoding the p110δ catalytic subunit or the p85α regulatory subunit of PI3Kδ, respectively (7). The most commonly detected variants in APDS patients are the E1021K substitution in p110δ, leading to APDS1, and the 434-475 deletion in p85α, resulting in APDS2 (8,9). Both mutations lead to gain-of-function (GOF) of PI3Kδ and overactivation of the downstream AKT/mTOR cascade (10–13). In the immune system, PI3Kδ GOF leads to skewed B cell populations towards a transitional phenotype, decreased numbers of naïve T cells and increased senescent T cells, resulting in impaired vaccine responses and overall immune dysfunction (11,14,15). Consequently, APDS patients present with recurrent infections, lymphoproliferation, autoinflammatory disease and lymphoma (8,9).

Although predominantly expressed in peripheral blood mononuclear cells (16), PI3Kδ is also detected in murine (12,17) and human (12,18) brain tissue. In the CNS, the PI3K/AKT/mTOR axis has been shown to play a crucial role in neuronal differentiation and migration (19,20). Accordingly, mutations along this pathway have been commonly associated with neurodevelopmental and neuropsychiatric disorders (21). Although few studies have focused on the specific role of distinct PI3K isoforms in the CNS, PI3Kδ has been proposed to regulate soma size, dendritic complexity and spine number (12,22,23), suggesting a contributing role towards neuronal morphology. Interestingly, 19-31% of APDS patients were reported to exhibit neurodevelopmental delay (8,9). However, the lack of systematic cognitive evaluation in these reports hinders the quantitative study of PI3Kδ on neurological function. Nonetheless, this putative behavioural role of PI3Kδ is further implied by the report of increased p110δ expression in a person with autism spectrum disorder (ASD) (24).

In this work, we investigated the role of PI3Kδ in motor and cognitive behaviour. We describe two related APDS patients and report, for the first time, a case of APDS-associated ASD. APDS patients presented with deficits in visuomotor integration, particularly in inhibition-recruiting and memory tasks, accentuated by the ASD phenotype in one of them. Additionally, we conducted an extensive battery of behavioural tests in an APDS mouse model (11), and show that p110δ^E1020K^ mice present with changes in locomotion, learning and repetitive behaviour patterning. Taken together, our data suggest that PI3Kδ GOF increases the risk of atypical behavioural development, supporting previous findings on the interplay between the CNS and the immune system.

## Results

### Immunological profile and neuropsychiatric manifestations of APDS patients

We present a 29-year-old male patient, P1, the second child of non-consanguineous parents of Caucasian descent (P2) (Table 1, 2) (15). Since the age of 9 months, P1 suffered from recurrent upper and lower respiratory tract infections and diarrhoea. At the age of 3.5 years, P1 was hospitalized for generalized lymphadenopathy due to EBV infection. He was subsequently diagnosed with common variable immunodeficiency, based on low serum IgG and IgA levels (with elevated IgM levels), and recurrent infectious complications for which intravenous immunoglobulin replacement therapy was initiated. At the age of 7, P1 developed auto-immune complications, including cutaneous manifestations, fever, arthritis, anaemia, thrombocytopenia and hepatosplenomegaly, with positive antinuclear antibody and anti-dsDNA titres, described as systemic lupus erythematosus (SLE)-like disease, for which immunosuppressive therapy was initiated. Other complications included liver cirrhosis due to auto-immune hepatitis with portal hypertension, requiring liver transplantation in December 2020. At age 22, genetic testing revealed a c.3061 G>A mutation in the *PIK3CD* gene, resulting in an E1021K substitution and APDS1 (25,26).

**Table 1:** P1 and P2 clinical characteristics. SLE, systemic lupus erythematosus; ↓, decreased compared to control age-matched range.

|  |  |  |
| --- | --- | --- |
| Sex | Male (P1) | Female (P2) |
| Age (diagnosis) | 29 (3.5 years) | 56 (childhood) |
| Mutation | E1021K | E1021K |
| Response to immunization | ↓ *S. pneumonia* (polysaccharide response) | ↓ *Influenza* type A and type B |
| Hepato/spleno megaly | Splenomegaly | Absent |
| Cytopenia | Leucopenia, thrombocytopenia | None |
| CT-chest results | Air trapping, no bronchiectasis | Bronchiectasis |
| Hematological malignancy | No | No |
| Other comorbidities | Psychomotor developmental delay<br>SLE-like auto-immune disease<br>Recurrent EBV infections<br>Autoimmune hepatitis with liver cirrhosis and portal hypertension | None |
| Ig therapy | Intravenous Ig replacement therapy: 35 g, every 3 weeks | Intravenous Ig replacement therapy: 15 g, every 4 weeks |
| Other relevant treatments | Prednisone: 10 mg, once daily<br>Mycophenolate mofetil: 500 mg, twice daily<br>Hydroxychloroquine: 200 mg, once daily<br>Trimethoprim/sulfamethoxazole: 480 mg, once daily | No immunosuppressive medication<br>No prophylactic antibiotics |

**Table 2:** P1 and P2 immunological findings. Abs, absolute numbers; ↓, decreased compared to control age-matched range.

| Patient | P1 | P2 |
| --- | --- | --- |
| Ig at diagnosis<br>(g/L; range (102)) | **IgG** 0.46 (4.0-11.0), **IgA** 0.45 (0.1-1.6),<br>**IgM** 3.24 (0.5-1.8) | *Unknown* |
| T/B/NK cells at diagnosis<br>(absx10<sup>9</sup>/L; range (103)) | **T cells** 3.66 (0.9-4.5), **CD4<sup>+</sup> T cells** 0.51<br>(0.5-2.4), **CD8<sup>+</sup> T cells** 3.15 (0.3-1.6), **B**<br>**cells** 0.26 (0.2-2.1), **NK cells** 0.26 (0.1-<br>1.0) | *Unknown* |
| T/B/NK cells<br>(absx10<sup>9</sup>/L; range (103))<br>[B cell subsets in abs, cells/ul] | **Age 22**<br>**CD3<sup>+</sup> T cells** 0.23 (0.7-2.1); **CD4<sup>+</sup> T**<br>**cells** 0.1 (0.3-1.4): Naïve<br>(CD4<sup>+</sup>/CD27<sup>+</sup>/CD45RA<sup>+</sup>) 14.4%,<br>Memory (CD4<sup>+</sup>/CD27<sup>+</sup>CD45RA<sup>-</sup>)<br>83.6%, Effector Memory<br>(CD4<sup>+</sup>/CD27<sup>+</sup>/CD45RA<sup>+/-</sup>) 2.0%; **CD8<sup>+</sup>**<br>**T cells** 0.1 (0.2-1.2): Naïve<br>(CD8<sup>+</sup>CD27<sup>+</sup>CD45RA<sup>+</sup>) 37.7%, Memory<br>(CD8<sup>+</sup>/CD27<sup>+</sup>/CD45RA<sup>-</sup>) 33.2%,<br>Effector Memory (CD8<sup>+</sup>/CD27<sup>-</sup><br>/CD45RA<sup>+/-</sup>) 29.1%; **CD19<sup>+</sup> B cells**<br>0.01 (0.1-0.5): Naïve (IgD<sup>+</sup>/CD27<sup>-</sup>) 7<br>(57-447), Marginal Zone/Natural<br>effector (IgD<sup>+</sup>/CD27<sup>+</sup>) 1 (9-88),<br>Memory (IgD<sup>-</sup>/CD27<sup>+</sup>) 1 (13-122) [IgM<sup>+</sup><br>48% (4-37), IgM<sup>-</sup> 52%]; **CD16.56<sup>+</sup>CD3<sup>-</sup>**<br>**NK cells** 0.04 (0.09-0.6) | **Age 43**<br>**CD3<sup>+</sup> T cells** 0.53 (0.7-2.1)<br>**CD19<sup>+</sup> B cells** 0.09 (0.1-0.5):<br>**CD16.56<sup>+</sup>CD3<sup>-</sup> NK cells** 0.15 (0.09-0.6) |
| Ig (g/L; range (102)) | **Age 28**<br>**IgG** 12.0 (6.0-12.3), **IgA** 0.43 (0.3-2.0),<br>**IgM** 3.24 (0.5-2.0) | **Age 55**<br>**IgG** 16.6 (6.0-12.3), **IgA** 1.49 (0.3-2.0),<br>**IgM** 4.75 (0.5-2.0) |

Besides this immunological phenotype, we also observed neuropsychological deficits in P1. Psychomotor developmental delay was present, as the patient started walking at the age of 2 and speaking at age 2.5. At age 6, ASD was considered and P1 was referred to special needs education. At the age of 9 years, intelligence quotient testing indicated a score of 80. Moreover, P1 showed persistent deficits in social interaction, motor function and a distinct fascination for watches, calendars and dates. P1 was diagnosed with pervasive developmental disorder not otherwise specified at age 10, and re-evaluation in 2020 confirmed the diagnosis of ASD based on psychiatric examination and on the autism-spectrum quotient (27). To date, P1 requires assistance with tying shoelaces and buttoning his shirts.

Patient 2 (P2), who has been previously described (15), is a non-consanguineous parent from P1. Genetic testing revealed a c.3061 G>A mutation in the *PIK3CD* gene, resulting in the E1021K substitution, which was also found in P1. P2 suffered from recurrent upper and lower respiratory tract infections since childhood and was diagnosed with an IgG2 and IgG4 subclass deficiency. She then commenced immunoglobulin replacement therapy and has been on intravenous treatment since. A recent CT-scan showed bronchiectasis. There have been no signs of hepatosplenomegaly nor lymphadenopathy. Currently, her clinical phenotype is relatively mild, with no recurrence of severe infections, no auto-immune complications, no inflammatory disease and no haematological malignancy. She was never diagnosed with a neurodevelopmental condition.

### APDS patients present with deficits in visuomotor integration

Previous clinical descriptions of APDS reported the presence of cognitive impairment, developmental delay or speech delay in a number of patients (8,9). Given the formal diagnosis of ASD in P1, and its association with attention and motor performance (28–30), we conducted a series of tests to evaluate visuomotor performance in both patients (Fig. 1a-c).

**Figure 1:**
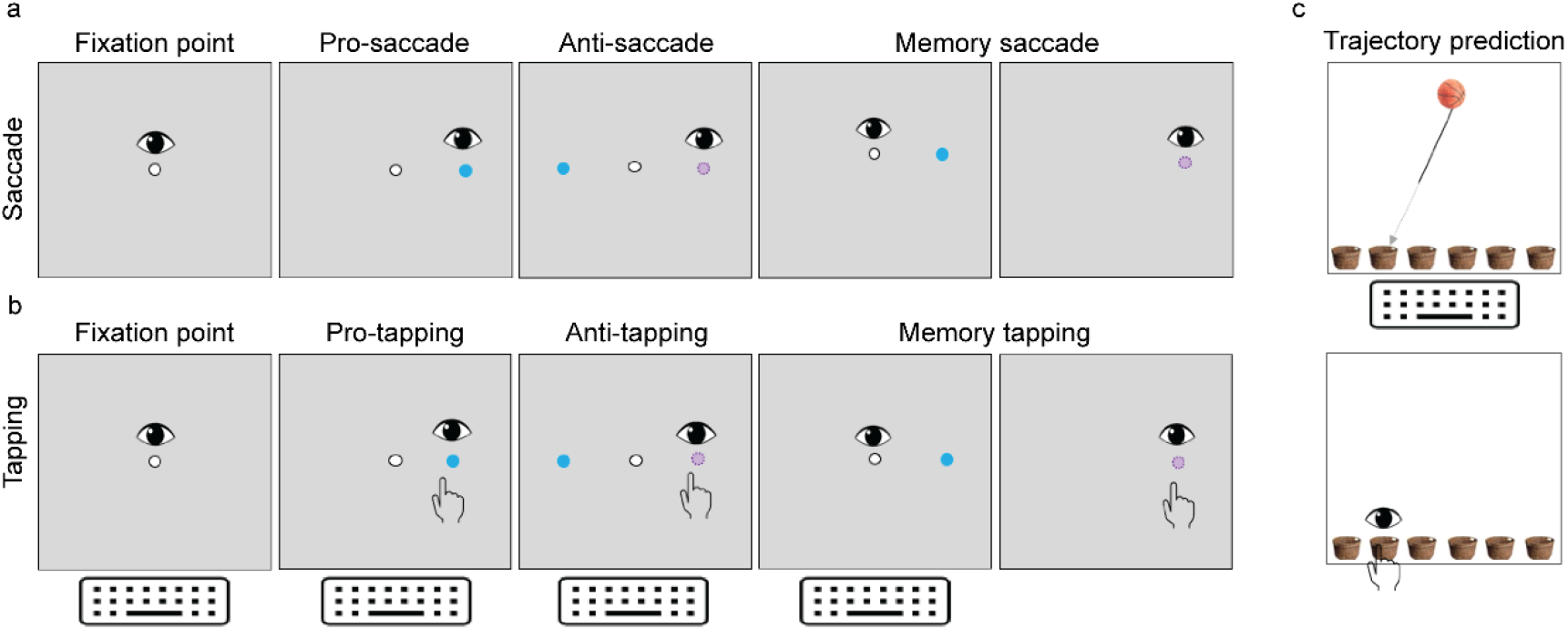
Visuomotor assessment tasks. Visual representation of the saccade (a), tapping (b) and trajectory prediction (c) tasks performed by patients P1 and P2, and respective age-matched controls. Pro-tasks involved the execution of reflexive saccades (and tapping) towards a newly appeared target while anti-tasks required a saccade execution (and tapping) to the opposite side of the new target. In memory tasks, subjects waited for target omission to perform a saccade (and tapping). Trajectory prediction tasks involved the execution of a saccade and tapping towards the basket in which a moving ball would be expected to fall. The number and age distribution of control participants per task can be found in Table 3.

Visual reflexive behaviour, primarily driven by parietal eye field and brainstem functions (31,32), was intact in both patients, with performance in the pro-saccade test equal between P1, P2 and their respective age-matched controls (performance: 100% for all groups) (Fig. 2a). While pro-tapping performance was also similar for all cohorts (performance: 100% for all groups), P1 exhibited increased hand latency compared to the other groups. Specifically, P1 average latency was over 4.5 SD higher than the age-matched control group (C1) (P1 = 463 ms, C1 = 391 ± 16 ms, P2 = 400 ms, C2 = 403 ± 33 ms) (Fig. 2a, c).

**Table 3:** Characteristics of controls. Age-matched individuals were tested in visuomotor assessment tasks and used as controls for patient 1 or patient 2. * for the trajectory prediction task, only one control group was used.

| Visuomotor assessment | Control 1 |  |  | Control 2 |  |  |
| --- | --- | --- | --- | --- | --- | --- |
|  | Age<br>(average, years) | SD | N | Age<br>(average, years) | SD | N |
| Pro-saccade | 24,82 | 5,04 | 12 | 54,00 | 13,39 | 11 |
| Anti-saccade | 25,00 | 3,35 | 14 | 57,70 | 8,66 | 17 |
| Memory saccade | 24,84 | 5,28 | 11 | 45,70 | 7,27 | 10 |
| Pro-tapping | 24,76 | 3,88 | 10 | 56,80 | 13,11 | 10 |
| Anti-tapping | 24,90 | 7,06 | 10 | 55,96 | 7,68 | 28 |
| Memory tapping | 25,47 | 5,67 | 10 | 53,62 | 6,31 | 16 |
| Trajectory prediction* | 38,30 | 8,78 | 10 | NA | NA | NA |

**Figure 2:**
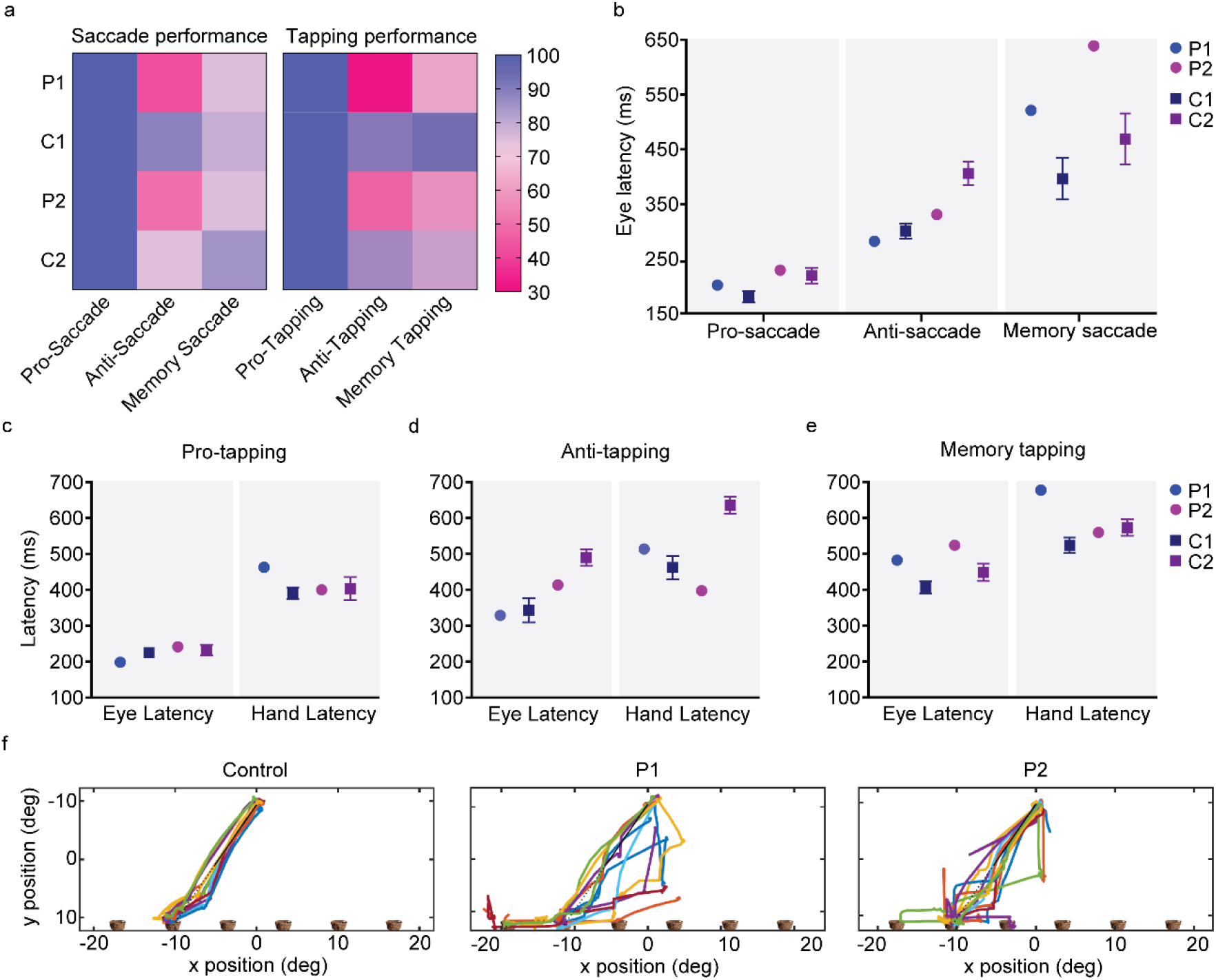
APDS patients present with intact reflexive saccades but altered integration. a) Performance in the saccade and tapping tasks is presented as percentage of correct trials. Eye latency for the saccade tasks (b), and eye and hand latency for the tapping tasks (c-e) are presented in ms. f) Representative traces of the eye trajectories performed towards one basket, during the trajectory prediction task. P1, patient 1, P2, patient 2; C1, age-matched controls for patient 1, C2, age-matched controls for patient 2.

To understand whether this increased hand latency was due to a motor impairment or rather a consequence of increased task complexity, motor command and execution were tested in the trajectory prediction test (Fig. 1c). Both patients exhibited similar latencies in decisive saccades towards the target basket, indicating that the task was correctly understood (P1 = 513 ms; P2 = 570 ms; C = 539 ms). Average hand latency was also similar for both P1 and P2 when compared to control groups, suggesting intact preparation and onset of motor response (P1 = 779 ms; P2 = 761 ms; C = 767 ms). However, while P2 exhibited similar performance to controls, P1 presented with a reduction in the percentage of correct trials (P1 = 78%, P2 = 100%, C = 96%) (Fig. 2f). Moreover, both APDS patients adopted a less systematic strategy to follow the ball’s trajectory compared to controls, exhibiting less goal-directed scan paths and more irregular eye gaze (Fig. 2f). These data suggest that, while preparation and onset of motor responses appear to be intact in both patients, increased task speed and complexity likely impairs integration, particularly in P1.

We next tested volitional inhibitory behaviour using the anti-saccade and anti-tapping tests (Fig.1a, b). Both tests require a suppression of reflexive pro-saccades and engage a complex network of brain regions, including dorsolateral prefrontal cortex, frontal eye fields, and supplementary eye fields, basal ganglia, superior colliculus and cerebellum (33–35). The anti-saccade task has been used to characterize cognitive impairments in patients with schizophrenia (36,37), dementia (38), Parkinson’s disease (39) and cerebellar atrophies (40). In the anti-saccade and anti-tapping tests, both APDS patients underperformed controls (anti-saccade performance: P1 = 43%, C1 = 89%, P2 = 50%, C2 = 74%; anti-tapping performance: P1 = 31%, C1 = 88%, P2 = 50%, C2 = 83%) (Fig. 2a). While patient eye latency was faster when compared to respective controls (P1 = 329 ms, C1 = 344 ± 33 ms, P2 = 414 ms, C2 = 490 ± 23 ms), indicative of frontal inhibition deficits (41), P1 hand latency was increased during tapping (P1 = 514 ms vs C1 = 462 ± 33 ms; 1.6 SD difference). P2 presented delayed hand execution time (time between screen bar release to target) (P2 = 992ms vs C2 = 132 ± 35ms; > 24.5 SD difference) combined with faster hand latency (P2 = 398 ms vs C2 = 636 ± 23 ms) (Fig. 2b, d). Together, our data show that P1 presents with movement integration deficits while P2, despite better performance to age-matched controls, exhibits delayed movement execution.

To further evaluate integration deficits, both patients performed a spatial memory task requiring both inhibition and memory retrieval (Fig. 1a, b). While performance in the memory-saccade task was similar for all groups (P1 = 75%, C1 = 79%, P2 = 759%, C2 = 85%) (Fig. 2a), both patients exhibited delayed eye latency (P1 = 483 ms, C1 = 406 ± 17 ms; > 4.5 SD difference, P2 = 524 ms, C2 = 449 ± 24 ms; > 3 SD difference) (Fig. 2b). In line with the anti-tapping task, P1 presented with increased hand latency, compared to controls (P1 = 678 ms vs C1 = 524 ± 21 ms; > 7 SD difference) whereas P2 exhibited severely delayed hand execution time (P2 = 919 ms vs C2=146 ± 39 ms; > 19.5 SD difference) (Fig. 2e). These results show that, while target location was remembered by both patients, in addition to the aforementioned motor integration deficits, recalling target position was delayed.

### PI3Kδ is expressed in adult mouse brain

Our patient data suggested that *PIK3CD* GOF increased the risk of neuropsychiatric dysfunction, supporting previous reports (8,9). To fully characterize the extent of neurological deficits and establish an animal model to test future pharmacological interventions, we resorted to a heterozygous mouse model of APDS (E1020K knock-in mouse, further referred to as “p110δ^E1020K^ mice”) (11), to explore the effects of *Pik3cd* GOF on behaviour.

Prior work in WT mice with a lacZ-*p110δ* reporter indicated the presence of p110*δ* in adult brain, predominantly in the cortex and hippocampus (17). Supporting these results, we detected an 110 kDa band in both WT and p110δ^E1020K^ brain tissue (Supplementary Fig. 1). p110δ was highly expressed in the spleen, as expected due to abundant B cell populations (42). In the brain, we found lower expression levels of p110δ, primarily detected in the cortex, hippocampus and olfactory bulbs (Supplementary Fig. 1).

### p110δ^E1020K^ *mice exhibit intact gross motor skills but altered locomotion pattern*

Having confirmed the expression of PI3Kδ in the brain, we proceeded with behaviour testing. We first assessed motor performance, which is found to be impaired in a number of patients with neurodevelopmental delay, particularly ASD (43,44). Spontaneous locomotion was tested in the open-field arena (Fig. 3a). Both WT and p110δ^E1020K^ mice moved more during the first 10 mins of exploration (Supplementary Fig. 2a), with mean speed and distance travelled across the total 30 mins of testing similar between genotypes (speed: *t*(28) = 0.5494, *p* = 0.59; distance: *t*(28) = 1.234, *p* = 0.22) (Fig. 3b,c). PI3Kδ mutation also did not affect performance on the rotarod test (Fig. 3d) (main effect of genotype, *F*(1,28) = 0.1789, *p* = 0.68), indicating that p110δ^E1020K^ mice have no gross motor defects.

**Figure 3:**
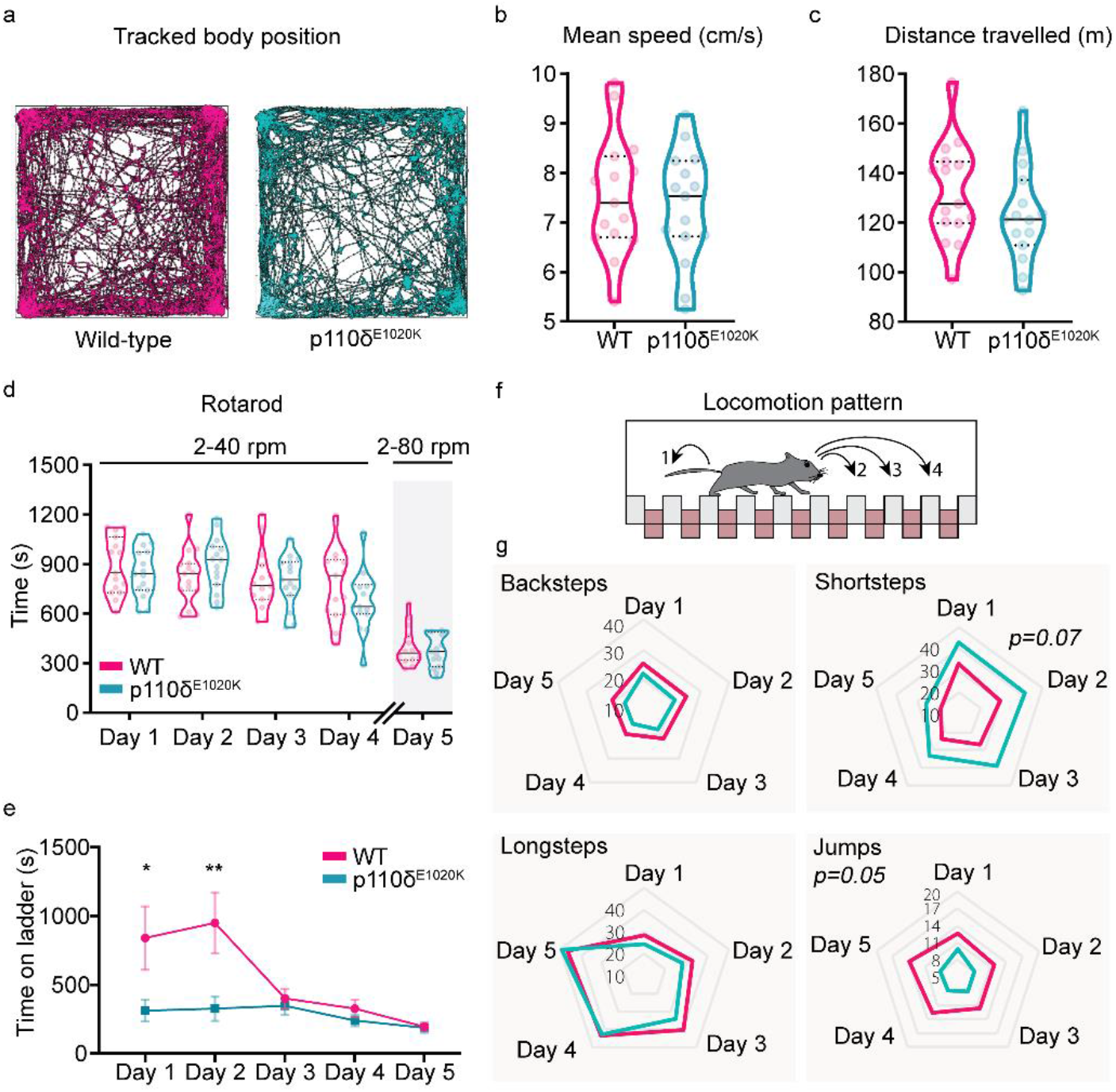
Slight fine locomotion impairments are caused by the murine E1020K mutation. a) Example of automatically tracked trajectories showing the body position of a WT and a p110δ^E1020K^ mouse during the 30 minutes of the OF task. b-c) Quantification of the mean speed (n = 15 per genotype) and total distance travelled (n = 15 WT and 14 p110δ^E1020K^) during the OF task, presented as median and quartiles (2-tailed t-test). d) The total time each mouse spent on the rotarod, over the course of 4 trials/day, is presented as median and quartiles (2-way repeated-measures ANOVA, n = 15 per genotype). On the last day, the maximum rod speed was increased to 80 rpm. e-g) The Erasmus ladder was used to investigate locomotion pattern. The average time each mouse spent on the ladder, across 42 daily trials, is presented in (e) (2-tailed Mann-Whitney; data presented as mean ± SEM). The distinct step types analysed are schematically represented in (f) and quantified in (g) (Mixed effects model; data is presented as daily mean percentage, n = 15 per genotype). * p≤0.05, ** p≤0.01, *** p≤0.001.

To investigate fine motor skills, mice were tested with the Erasmus ladder, a fully automated behavioural apparatus that allows detailed analysis and quantification of motor performance and learning in mice (45). p110δ^E1020K^ mice spent significantly less time crossing the ladder on the first two days of testing (Day 1: *U* = 60, p = 0.05; Day 2: *U* = 40, p = 0.01) (Fig. 3e). This was not prompted by a higher efficiency in crossing the ladder, as the percentage of missteps was similar for each day in both genotypes (main effect of genotype: *F*(1,28) = 1.786, p = 0.19) (Supplementary Fig. 2c). We next analysed the locomotion pattern on the ladder (Fig. 3f). Although WT and p110δ^E1020K^ had identical percentages of backsteps (*F*(1,28) = 2.784, p = 0.11) and longsteps (*F*(1,28) = 0.4735, p = 0.50), p110δ^E1020K^ mice displayed a tendency to use a higher percentage of short steps (*F*(1,28) = 3.469, p = 0.07) and used fewer jumps (F(1,28) = 4.112, p = 0.05) to cross the ladder (Fig. 3g). These pattern changes were independent of weight as this progressed similarly between groups (Supplementary Fig. 3).

Together, these results indicate that PI3Kδ GOF mutation has no impact on gross motor function, but contributes to changes in fine locomotor skills that result in the adoption of a different locomotion strategy by mutant mice. This is in line with the findings in our patients, who do not present with gross motor function impairments either, but do present with fine-motor movement impairments.

### p110δ^E1020K^ *mice show altered patterns of repetitive behaviour independent of anxiety-like measures*

PIDs predispose patients to an increased prevalence of mood disorders (46), as does the presence of developmental delays (47,48). To further investigate anxiety-like behaviour in p110δ^E1020K^ mutants, we tested mice in the open-field (OF) and elevated-plus maze (EPM) tests.

In the OF, we found no evidence of increased anxiety-like behaviour in p110δ^E1020K^ mice, as both genotypes spent comparable time in the inner and outer areas of the arena (In: U = 67, p = 0.10; Out: U = 103, p = 0.71), as well as in the corners (U = 111, p = 97) (Fig. 4a,b, Supplementary Fig.2b). There was also no effect of genotype in the EPM regarding the time spent on the different arms of the maze (F(1,28) = 0.10, p = 0.75) or the number of transitions between arms (F(1,28) = 3.18, p = 0.09) (Fig. 4c,d). These data indicate that p110δ^E1020K^ mice do not exhibit increased anxiety despite their immunological phenotype (11).

**Figure 4:**
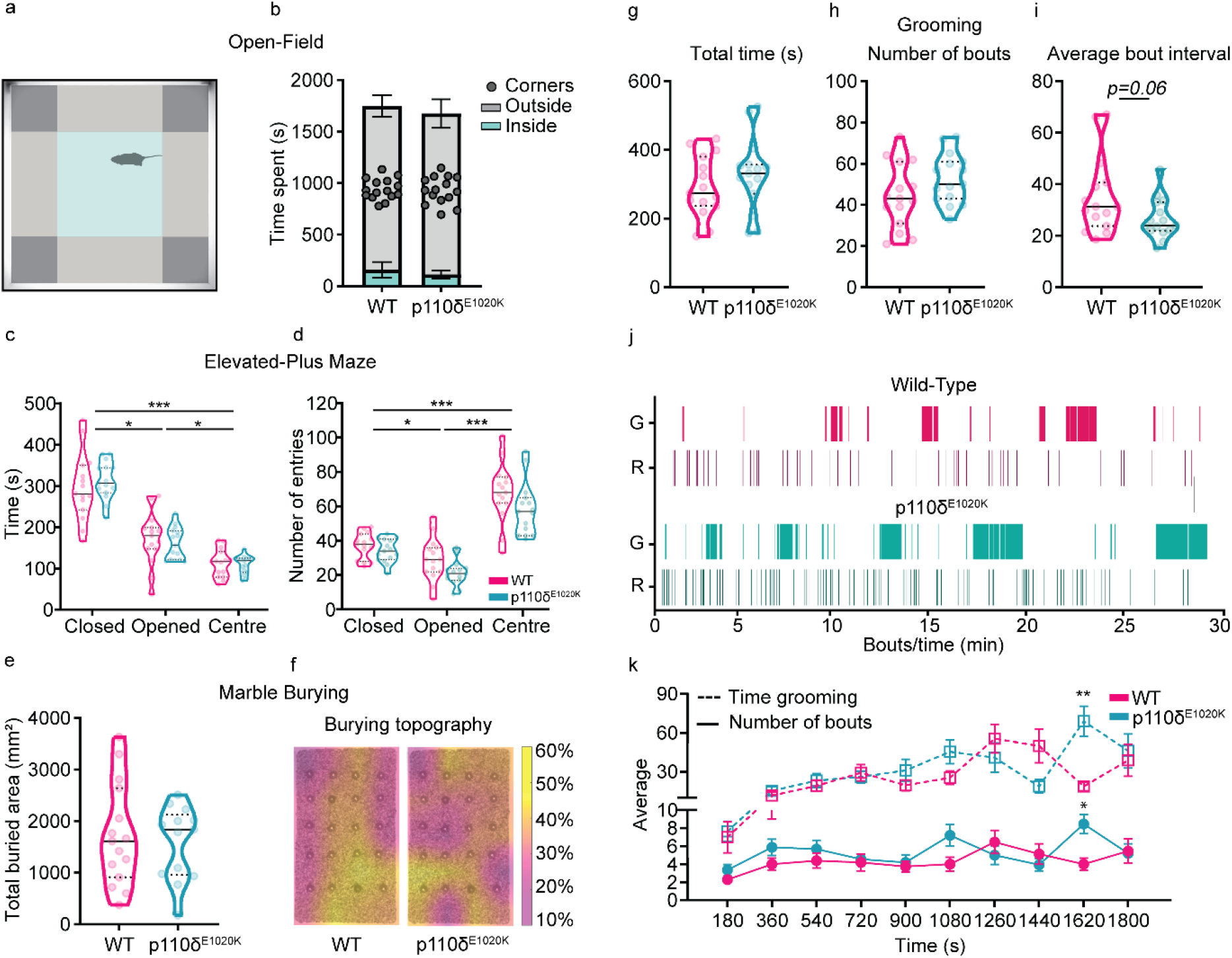
p110δ^E1020K^ mice exhibit subtle changes in burying and grooming patterns. a) Representation of the OF arena parcellation into corner, outside and inside areas. b) Total time spent on each OF area (2-tailed Mann-Whitney; data presented as mean ± SD). c-d) Total time spent and number of entries performed in each EPM area (2-way repeated-measures ANOVA; data are presented as median and quartiles). e) Total marble area buried during the MB task (2-tailed t-test). f) Marble disposition before the task, superimposed with the average percentage of buried area per marble (n = 15 WT and n = 13 p110δ^E1020K^). g-i) Quantification of the total time spent grooming (g), total number of grooming bouts (h) and the average time interval between grooming bouts (i), during the grooming assay (2-tailed t-test; data presented as median and quartiles). j) Representative plot depicting grooming and rearing events for one mouse of each genotype. k) Time-binned plot with the average time spent grooming (dashed line; 2-way repeated-measures ANOVA) and the average number of grooming bouts (full line; mixed effects model) (data are presented as mean ± SEM). G, grooming, R, rearing; * p≤0.05, ** p≤0.01, *** p≤0.001, n = 15 mice per genotype, expect for e) and f) (see above).

We next explored the presence of repetitive behaviours, a common comorbidity of neurodevelopmental delays (49). Using the marble burying task (Supplementary Fig. 4a,b), we first measured the total area buried by each mouse and found this to be similar between genotypes (area buried: t(26) = 0.43, p = 0.67; number of buried marbles: t(26) = 0.00, p > 0.99) (Fig. 4e, Supplementary Fig. 4c). When the location of the buried marbles was mapped, we found that WT mice preferably buried marbles in the bottom right corner and centre, while p110δ^E1020K^ mice favoured areas close to the walls of the arena (Fig. 4f), indicating increased thigmotactic behaviour.

As the previous results suggested the presence of a distinct repetitive behaviour pattern, we further addressed this using the grooming assay. The total time spent grooming was similar between groups (t(28) = 0.96, p = 0.34) (Fig. 4g), as was the total number of grooming bouts (t(28) = 1.65, p = 0.11) (Fig. 4h) and latency to initiate grooming behaviour (t(28)=0.25, p = 0.81) (Supplementary Fig. 4d). We found a tendency for the average time interval between grooming bouts to be smaller in p110δ^E1020K^ mice (t(28) = 1.98, p = 0.06) (Fig. 4i, j), further suggesting a difference in behaviour pattern between groups. Indeed, a significant interaction between genotype and short and long grooming bouts (genotype x type of bout: F(1,28) = 5.31, p = 0.03) revealed that p110δ^E1020K^ mice exhibited a higher prevalence of short bouts compared to WT, while the opposite was observed for the long bouts (short bouts: 13.29% in WT vs 19.46% in p110δ^E1020K^; long bouts: 86.71% in WT vs 80.54% in p110δ^E1020K^) (Supplementary Fig. 4e). Furthermore, when the number of grooming bouts was analysed over time, a tendency for increased bout number over time was seen in p110δ^E1020K^ mice towards the end of the assay (genotype x time: F(9,251)=2.15, p = 0.03) (Fig. 4k, bottom curve). A significant time x genotype interaction was also found for the 3 minute-binned time spent grooming (F(9,273) = 3.51, p = 0.0004) (Fig. 4k, top curve), further supporting the presence of an altered grooming pattern in p110δ^E1020K^ mice. Taken together, these data indicate that, despite the absence of increased anxiety-like behaviour, p110δ^E1020K^ mice present with subtle alterations in the pattern of repetitive behaviour.

### p110δ^E1020K^ mice exhibit changes in associative response

Given the cognitive impairments and learning difficulties presented by some APDS patients (7,9), including P1, we investigated associative and spatial learning in the p110δ^E1020K^ mice.

First, we quantified learning using the Erasmus ladder. During the Erasmus ladder task, two stimuli are presented. Initially, a light turns on inside the goal box. Next, an air stream encourages the mouse to enter the ladder (45) (Fig. 5a). Considering the exit frequency for each stimulus, we found that both genotypes responded similarly to stimuli in the first sessions of the task. For sessions 3 and 5, p110δ^E1020K^ mice left the goal box less frequently with the air stimulus than WT (session 3: U = 61, p = 0.03; session 5: U = 55.5, p = 0.02) (Fig. 5b), while increasing box exits after light presentation in later sessions (session 4: U = 63.5, p = 0.07; session 5: U = 63.5, p = 0.04) (Fig. 5c,e). Increased light exit frequency could be representative of increased readiness or impulsivity to leave the box, interrupting the pre-stimulus waiting period. From testing days 1 to 4, both genotypes left the box before cue presentation with similar frequencies (Fig. 5d). On day 5, this frequency was increased in p110δ^E1020K^ mice (session 5: U = 60.5, p = 0.05). As expected, there was a positive association between leaving before cue and the light/air exit ratio (WT: ⍴ = 0.49, p < 0.0001; p110δ^E1020K^: ⍴ = 0.63, p < 0.0001). Least squares fitting demonstrated that the response of the two genotypes to the stimuli was significantly different (F(2,139)=3.906, p = 0.02; WT: y = 4.9x+3.8; p110δ^E1020K^: y = 2.3x+4.5) (Fig. 5f).

**Figure 5:**
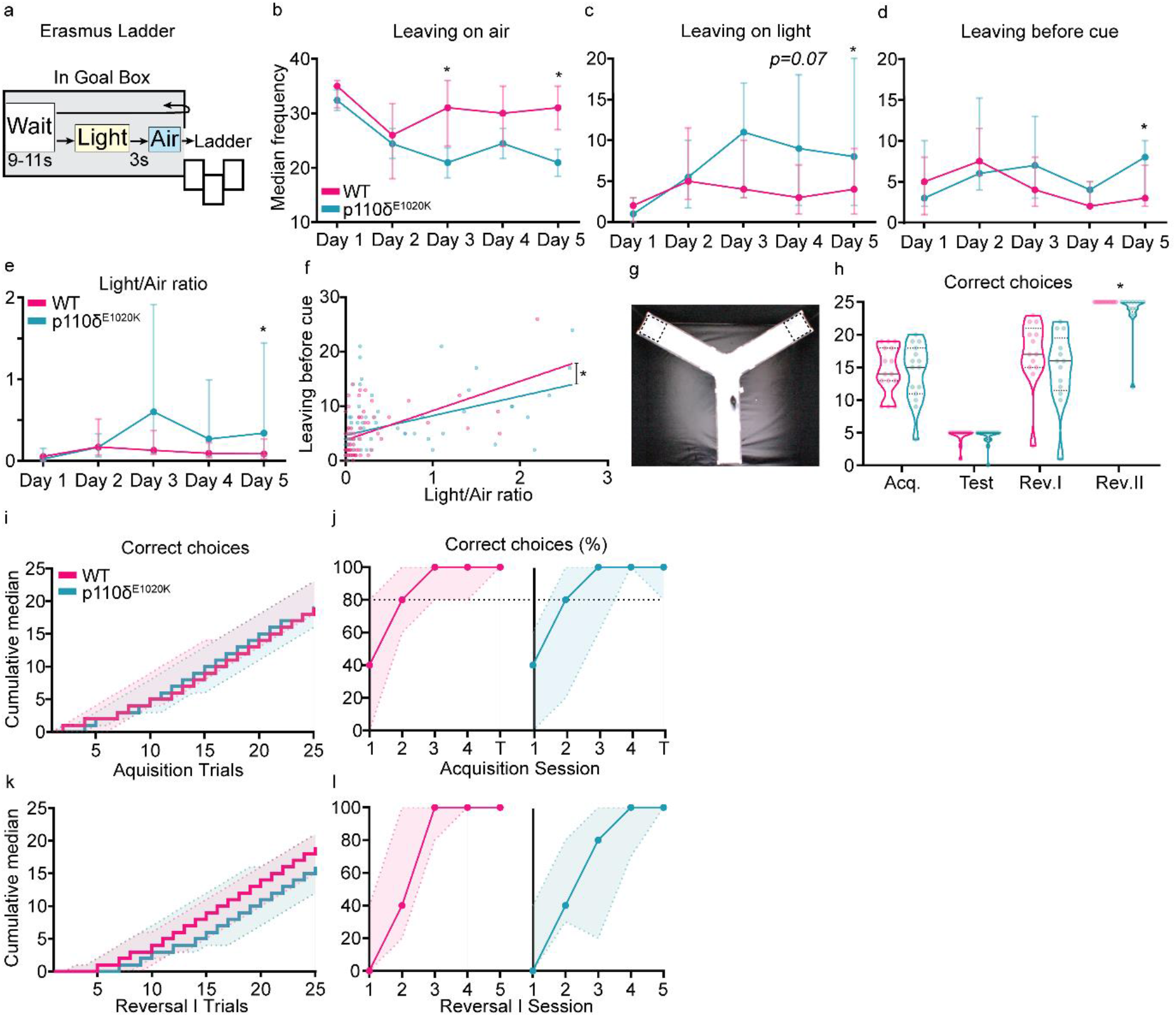
Stimulus-dependent ladder exit and reversal learning are mildly affected in p110δ^E1020K^ mice. a) Schematic of the Erasmus ladder goal-box with the time intervals between stimuli. b-d) Number of times individual mice left the goal-box with the air stimulus (b), the light stimulus (c), or before light cue presentation (d). e) ratio between light and air exits. f) Best-fit regression model between the data points used to plot (d) and (e). g) Picture of the Y-Maze, with dashed squares representing the possible locations for the hidden platform, either on the right or left arm of the apparatus. h) Total number of correct arm choices for both genotypes, during each phase of the Y-Maze (data presented as median with interquartile range). i) Step function with the cumulative median and interquartile range for the number of correct arm choices during all acquisition and test trials. j) Percentage of correct arm choices for each genotype over the four days of acquisition and the day of test (data presented as median with interquartile range) k-l) Similar to (i) and (j) but for the reversal I phase. 2-tailed Mann-Whitney, except for f). * p≤0.05, n = 15 mice per genotype, except for reversal phases where n = 13.

To further explore learning behaviour, we used the water Y-maze (Fig. 5g), a test often used to study repetitive behaviour and cognitive flexibility in ASD-mouse models (50–52). Similar to the previous OF and Rotarod results, we found no evidence of motor dysfunction, with both genotypes swimming similar distances and at comparable speeds during the habituation phase (distance: U = 67, p = 0.22; speed: U = 77, p = 0.23) (Supplementary Fig. 5a,b). During the acquisition and test phases, both WT and p110δ^E1020K^ mice learned the platform location, and there was no difference in the number of correct arm choices made by each genotype (Fig. 5i,j). When the location of the platform was reversed, p110δ^E1020K^ mice presented with a lower cumulative median of correct choices per trial, taking longer to perform the task correctly (Fig. 5k). No significant differences were found in the total number of correct choices per session (Fig. 5l). Similar results were obtained regarding the reversal II phase (Supplementary Fig. 5d,e). However, in this phase, errors in platform arm choice were only performed by p110δ^E1020K^ mice (U = 45.5, p = 0.01) (Supplementary Fig. 5e). Taken together, these results indicate that p110δ^E1020K^ mice present with mild deficits in paired-stimulus learning and reversal learning.

### p110δ^E1020K^ mice display intact social interaction behaviour

Atypical development of social skills and interactions is a common component of neuropsychiatric conditions, particularly of those with ASD comorbidity (53,54). Therefore, we sought to evaluate the performance of p110δ^E1020K^ mice in a social interaction paradigm (55).

During baseline exploration of the three-chamber apparatus, when no social stimulus was presented, both genotypes displayed a similar ambulatory behaviour across all chambers (F(1,28) = 0.5376, p = 0.47) (Supplementary Fig. 6a,c). p110δ^E1020K^ mice displayed slightly altered exploratory behaviour, with a tendency for centre crossing avoidance (genotype x chamber: F(2,55) = 2.988, p= 0.06; WT mean centre transitions = 46.57 vs p110δ^E1020K^ mean centre transitions = 38.40) (Supplementary Fig. 6d). During the test phase, a novel mouse was introduced to the arena (Fig. 6a). Both WT and p110δ^E1020K^ mice spent more time in the chamber where the novel mouse was located (main effect of chamber: F(1.911,79.30) = 87.71, p < 0.0001; main effect of genotype: F(1,83) = 0.0006, p = 0.98) (Fig. 6b), increasing the time spent in this chamber compared to their correspondent baseline values (main effect of phase: F(1,28) = 98.74, p < 0.0001; main effect of genotype: F(1,28) = 0.2441, p = 0.63) (Fig. 6c). Similar to what was found for the baseline exploration period, the avoidance of central area crossings in p110δ^E1020K^ mice persisted in the test phase (genotype x chamber: F(2,54) = 5.423, p = 0.01; WT mean centre transitions = 27.21 vs p110δ^E1020K^ mean centre transitions = 23.64) (Supplementary Fig. 6e).

**Figure 6:**
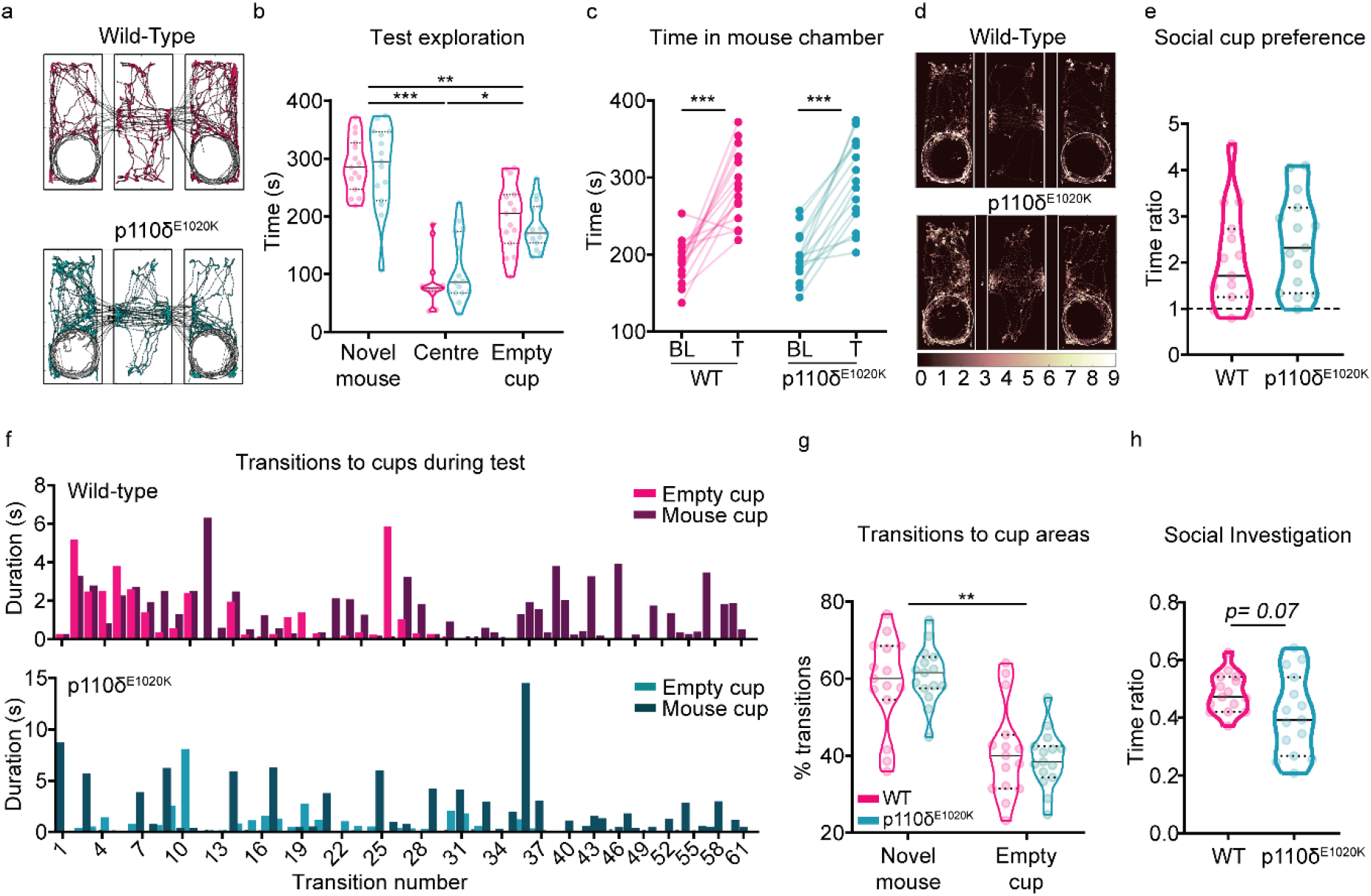
Social behaviour is largely preserved in p110δ^E1020K^ mice. a) Example of automatically tracked body positions during test phase (novel mouse on the left). b) Total time individual mice spent in each chamber of the apparatus during test phase (mixed effects model; data presented as median with interquartile range). c) Before and after plot of the total time each individual mouse spent on the novel mouse chamber during baseline (BL) and test (T) (2-way repeated-measures ANOVA). d) Body position heatmap depicting position frequency per 2.5mm bins (novel mouse on the left). e) Median and quartiles with the ratio between the time each individual mouse spent near the social cup over the time it spent near the empty cup (2-tailed t-test). f) Representative plot with the duration, in seconds, of each transition into the empty (light bars) or novel mouse (dark bars) cup area. g) Median and quartiles with the percentage of transitions each individual mouse made to the novel mouse or empty cup (2-way repeated-measures ANOVA); h) Median and quartiles of the ratio between the time individual mice spent exploring the novel mouse cup over the time spent in the whole novel mouse chamber (2-tailed t-test). * p≤0.05, ** p ≤ 0.01, *** p≤0.001. n = 15 mice per genotype.

Focusing on the region of interest defined around the empty cup and the cup with the novel mouse, both genotypes demonstrated a comparable preference for interacting with the cup where the social stimulus was located (t(28) = 0.99, p = 0.33), spending approximately twice the time exploring this cup compared to the empty cup (Fig. 6d,e). This preference for social cup exploration was also accompanied by an increased number of transitions into the novel mouse cup area (main effect of cup: F(1,28) = 29.04, p < 0.0001) (Fig. 6f, g). For both genotypes, the time spent exploring the novel social stimulus progressively decreased over the course of the task (main effect of time: F(1.328,118.8) = 5.714, p = 0.0002; main effect of genotype: F(1,28) = 0.3303, p = 0.57) (Supplementary Fig. 6b). Finally, when social investigation preference was analysed, p110δ^E1020K^ mice exhibited a tendency to spend a lower proportion of their time in the novel mouse chamber in the proximity of the cup, although this did not reach the statistical significance threshold (t(28) = 1.877, p = 0.07) (Fig.6h). Altogether, these data indicate that, despite a slight centre avoidance phenotype, p110δ^E1020K^ mice prefer the social stimulus over the asocial one, exhibiting an unaffected social phenotype.

### Motor, learning and repetitive behaviours best discriminate WT and p110δ^E1020K^ mouse populations

The analysis of independent readouts for each behaviour revealed a number of discrete changes in the behavioural pattern of p110δ^E1020K^ mice. Nonetheless, behaviour is a dynamic process where small stereotyped modules are often grouped or combined into larger representations that underlie each individual’s phenotype (56,57). To better understand the most important contributors to the phenotype of p110δ^E1020K^ mice, we performed linear discriminant analysis (LDA) on all behavioural variables measured (58,59). This type of analysis allows for encompassing individual differences across individuals and captures stable traits best separating the genotypes across many tests (59). We then selected the first two dimensions, LD1 and LD2 (Fig. 7a), and plotted the 10 best contributing components of each discriminant, as these are the variables that give the most information on group separation (Fig. 7b).

**Figure 7:**
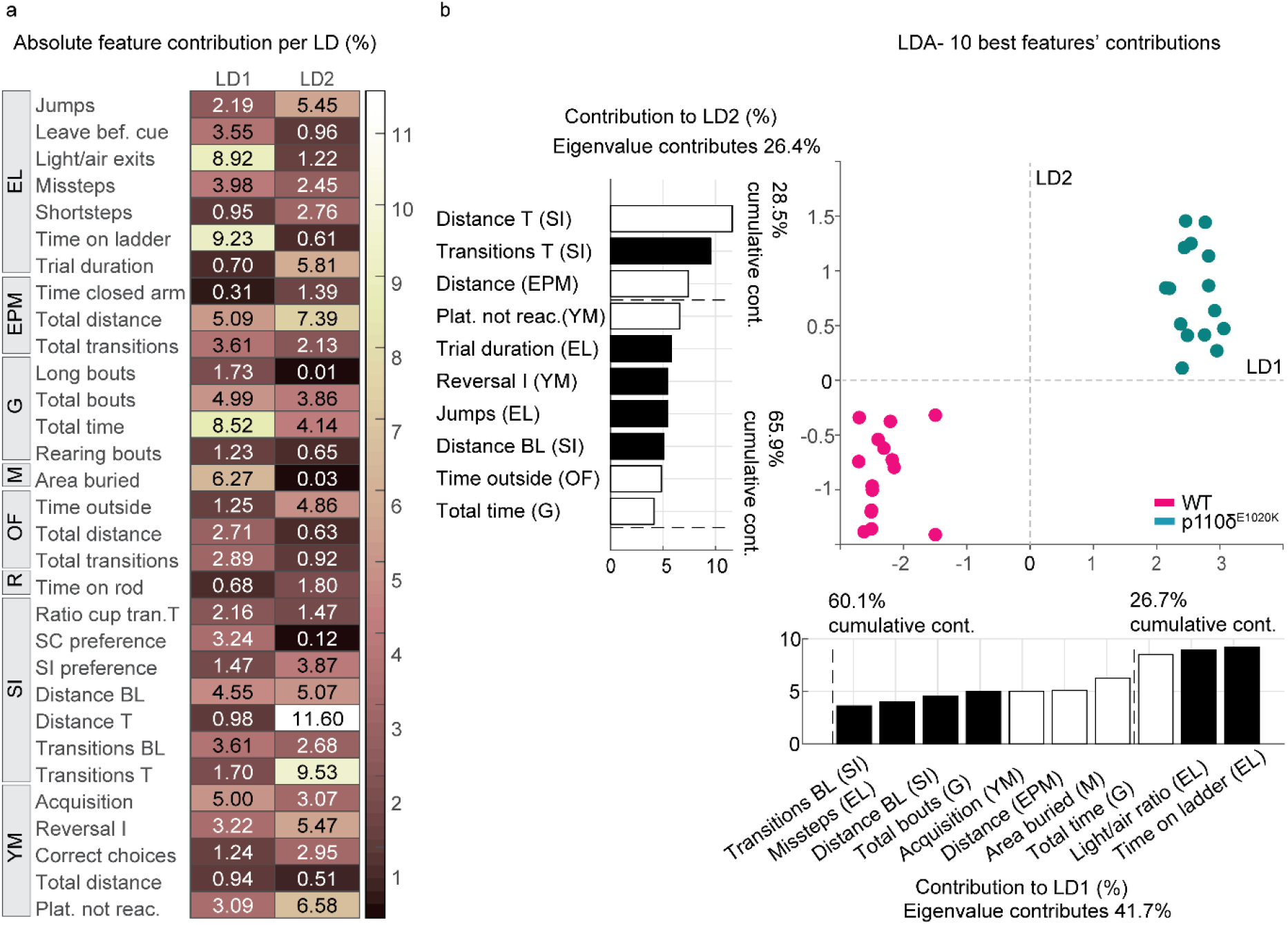
LD1 and LD2 features separate WT from p110δ^E1020K^ mice. a) Absolute contribution of each behavioural variable, in percentage, to linear discriminants 1 and 2, resulting from the LDA. b) LDA plot featuring the 10 best contributors to LD1 and LD2. Negative and positive contributions are represented by black and white bars, respectively. Each dot represents one mouse, with pink dots representing WT mice and green dots p110δ^E1020K^ mice. n = 15 mice per genotype. EL, Erasmus ladder, EPM, elevated-plus maze, G, grooming, M, marble burying, OF, open-field, R, rotarod, SI, social interaction, YM, water y-maze.

LDA of the behavioural data classified individual points into 2 non-overlapping classes, identifying the two genotypes. The 2 best LDs represent 68.1% of data variation, with LD1, which explains 41.7% of total data variation, creating a maximal separation between classes. Focusing on the greatest weights, motor and learning related variables (*time on ladder* and *light to air ratio*, respectively) contribute the most for group classification. The third feature, *total grooming time*, with an absolute contribution of 8.5%, indicates that additional group separation is achieved by the inclusion of repetitive behaviours in this discriminant. Further separation of the data along the vertical axis is provided by LD2, albeit with lower contributions (26.4%). This discriminant represents parameters predominantly influenced by locomotion-derived features. These include total distance travelled and transitions made during the test phase of the SI, and total distance travelled during the EPM. Altogether, these results indicate that LDA compiles and captures behavioural alterations in locomotor performance, learning and repetitive behaviours between WT and p110δ^E1020K^ mice, supporting the previously identified univariate analysis findings.

## Discussion

The study of the immune system in the regulation of neurodevelopment and in shaping subsequent behaviour is a rapidly emerging field, involving crosstalk in immunoneuropsychiatry and new integrative therapeutic approaches (60–62). In this work, we investigated neurologically-relevant behavioural features in APDS, a rare PID, using both patient data and a murine model. To our knowledge, this is the first study of APDS which specifically focuses on its behavioural component.

APDS patients exhibited changes in visuomotor responses, with P1 presenting with motor integration deficits, while both patients displayed decreased memory recall capacity. Additionally, P1 was also formally diagnosed with ASD, strengthening on previous more general reports describing neurodevelopmental delay as a comorbidity of APDS patients (8,9,13,63). In the p110δ^E1020K^ murine model, we detected more subtle phenotypic alterations. GOF mice presented with altered patterns of locomotion and repetitive behaviours, features reminiscent of symptoms found in individuals with ASD (49,64).

While our data supported a role of *PIK3CD* GOF on behaviour, the precise function of PI3Kδ in the brain remains elusive. In mice, p110δ has been found in brain and spinal cord, and proposed to have a role in neuronal morphology (12,17,22,23). Although the expression pattern of human *PIK3CD* follows a similar distribution as in mice (Allen Human Brain Atlas (2010)), reports of its non-immunological functions are scarce, with only a few studies implicating this isoform in schizophrenia and autism (18,24,65). The presence of behavioural deficits in adult mice combined with the low PI3Kδ expression in the brain, suggests that this isoform might have a predominant function during brain development rather than adulthood. Consistent with this hypothesis, recent studies found more PIK3CD transcripts in human foetal brain than in adult samples (18) and distributed expression of *PIK3CD* in the developing mouse brain (66). Combined with the fact that PI3Kδ lies upstream of the mTOR pathway, a signalling hub that is highly active during brain development and often found to be dysregulated in neurodevelopmental disorders (19,67,68), this expression suggests that *PIK3CD* plays a still unexplored role in the modulation of brain development, adding to the growing body of evidence pointing to a critical period for ASD development (52,69).

Despite its presence in the brain, PI3Kδ is predominantly expressed in leukocytes. This supports a putative modulatory role of the immune system on behaviour. Accordingly, a number of studies has now suggested a link between neurodevelopmental and immunological dysregulation. For example, in rodents, externally triggering a maternal immune response during pregnancy induces behavioural alterations in adult offspring. These include reduced cognitive flexibility and decreased social exploration, traits of an ASD-like phenotype (70–72). In humans, increased odds of neonatal infections were reported for children with ASD (73). Additionally, viral or bacterial infections during pregnancy were associated with an increased likelihood of ASD diagnosis (74,75), whereas increased ASD symptom severity was found in children with a maternal history of chronic immune activation (76). In APDS, family history of immunodeficiency is also estimated in 39% of patients (7), thus suggesting that PID might be an important predictor of neuropsychiatric load. Strengthening the hypothesis of an *immunological-behavioural phenotype* relationship is the presentation of the described patients, with P1 displaying increased immunodeficiency and visuomotor impairments, in addition to an ASD diagnosis, when compared to P2. Overall, the indirect links between immune and neuropsychiatric dysfunction indicates that immunological burden may be an important predictor for the development of atypical behaviour, not only in APDS, but also in other PID (77,78).

Interestingly, in individuals with ASD and ASD animal models, altered microbiota has been reported, with studies describing lower diversity profiles of colonizing microbiota in these groups (79–82). Highlighting a putative ASD-microbiota relationship, a recent follow-up study on microbial transfer therapy reported that ASD participants still scored 47% lower than baseline on the Childhood Autism Rating Scale and 35% lower on the Aberrant Behavior Checklist two years after trial conclusion, suggesting that microbiota regularization may improve autism-related scores on a longer term (83). Importantly, preclinical evidence suggests that PI3Kδ may play a role in microbiota regulation. Indeed, *Pik3cd*^E1020K/+^ mice were shown to exhibit increased antibody production and reactivity against autologous commensal bacteria (84), whereas mice with PI3Kδ loss-of-function develop colitis due to pathogenic T cell responses and altered IL-10 and IL-12p40 production (85,86). Thus, although human data are essential to further explore these hypotheses, given that 22% to 29% of APDS patients present with enteropathy (7), it would be important for future PI3Kδ studies to consider its possible role in microbiota homeostasis.

The interplay between the immune system and the brain is a current topic of rapid scientific discovery (87,88). Here, we show that a heterozygous mouse model of APDS displays mild behavioural alterations in addition to its immunological phenotype. ADPS patients showed high levels of heterogeneity when it came to behavioural and immunological symptoms. However, both P1 and P2 presented with sensorimotor deficits, a feature captured by the mouse model. Notably, the severity of the symptoms between P1 and P2 was reflected in the performance during the visuomotor tests. This is of interest to the APDS community, because such tests have previously been shown to accurately capture the features of early stage Alzheimer's disease (89), correlate with cognitive impairments in Parkinson’s disease (90), and serve as a tool to monitor the progression of both conditions (91). Further, due to their non-invasive nature, such tests are suitable to use even in very young children (92). In the future, we aim to further explore the correlation between immune system impairments, behavioural deficits and the outcome of the visuomotor deficits, on a larger APDS patient cohort, assessing the potential benefits of including this type of test batteries in the diagnostic pathway.

In addition to reinforcing the need for a multidisciplinary team assessing APDS patients, this study highlights the importance of increased monitoring of immunodeficient patients for the presence of neuropsychiatric comorbidities and describes a set of non-invasive tools that allow for such assessment. Additional studies on the function of PI3Kδ in the brain will be fundamental to understand its specific role in neurodevelopment and deepen our knowledge of the interactions between immunological burden and neuropsychiatric load.

## Methods

### Patients

We describe two APDS patients (Table 1,2). P1 is regularly followed (by VD) at the Primary Immunodeficiency Center of the Department of Internal Medicine, Division of Clinical Immunology, Erasmus MC (Rotterdam, The Netherlands); P2 is currently under treatment (by SMA) at the outpatient Department of Infectious Diseases of Leiden University Medical Center (Leiden, The Netherlands). Psychiatric assessment was performed (by NB) at the Erasmus MC and included the autism-spectrum quotient (27). Additional clinical history and data were obtained from medical notes (by OM, VD and SMA).

### Visuomotor coordination and memory assessment

An eye-hand coordination measurement setup was used to quantify the interactions between visual, ocular motor, and manual motor systems in both spatial and temporal domains. It consisted of a 21.5’’ touchscreen monitor (Wacom DTH-2242, Wacom Corporation, Japan), a remote infrared and screen-based eye-tracker (Tobii Pro X3, Tobii Corporation, Sweden) and a wired keyboard. The eye-tracker, positioned below the touchscreen, recorded eye movements at 120Hz and was connected via an external processing unit to a laptop (DELL Latitude 5590, Dell Technologies, Texas, United States) with an Intel Core i5-8350U processor, 256 GB SSD, and 16 GB internal RAM to warrant optimal performance and data quality (Pro, 2017). Eye movements with a speed > 50°/s were considered saccades. Manual responses were captured by sampling alternating presses and releases of the index finger from the dominant hand, between keyboard and touchscreen. After a short general instruction, each subject was instructed to sit straight in front of the touchscreen. Eye positions were calibrated at approximately 65 cm from the touchscreen using a standard calibration procedure. Next, 7 tasks, 3 eye tasks and 4 eye-hand tasks (Fig. 1), were presented on the touch screen in a fixed order (see below). Standard verbal instructions were given prior to each task and each subject was allowed a maximum of three practice trials. These instructions were also written on the screen (in Dutch). The starting position at each trial was fixating a central white dot and, in case of eye-hand tasks, also touching a blue bar at the bottom of the screen with the index finger for 2 seconds. The following 16 trials within each task had to be executed as fast and as accurately as possible. The following tasks were performed:

1. Pro-Saccade and 2. Pro-tapping: The subject had to fixate (pro-saccade) or touch (pro-tapping) a randomly appearing peripheral dot.
3. Anti-Saccade and 4. Anti-Tapping: The participant had to make an eye movement (anti-saccade) or an eye and hand movement (anti-tapping) in the opposite direction of a randomly appearing peripheral dot, at either 5, 10, 15 or 20 degrees of the horizontal direction.
5. Memory-Saccade and 6. Memory-Tapping: While fixating the central dot, a peripheral dot appeared for 50 ms at a random position. The subject had to fixate (memory-saccade) or touch (memory-tapping) the remembered peripheral dot location after the central dot disappeared.
7. Trajectory Prediction: A ball was dropped in the direction of one of six baskets. Halfway along the trajectory, the ball became invisible and the subject had to touch the basket in which the ball would have fallen.

### Mice procedures

Thirty 10 to 13 week-old wild-type (n = 15 WT) and heterozygous p110δ^E1020K^ (n = 15 p110δ^E1020K^) male mice were kindly provided by Dr. Klaus Okkenhaug (University of Cambridge, United Kingdom). These mice harbour an E1020K knock-in mutation in the *Pik3cd* gene expressed in all cells (11). Following arrival to the Erasmus MC, mice were acclimated to the facilities for two weeks. Mice were group-housed (3 to 4 mice per cage, mixed genotypes in the same cage), provided with food and water *ad libitum* and kept on a regular 12h light/dark cycle. After this acclimatization time, mice were handled by the experimenters for three days prior to experiment initiation. Before each experiment, mice were weighed (Supplementary Figure 3) and habituated to the testing room for at least 1 h. Experimenters were blinded to the genotype of each mouse.

When all behavioural experiments were completed, brain tissue was collected. Mice were injected with an overdose of pentobarbital, transcardially perfused with 0,9% NaCl, and the brain dissected. Tissue was flash frozen and kept at −80°C until used.

### Genotyping

Mice were genotyped by amplifying the *Pik3cd* locus from mouse ear DNA using the forward E1020KrecF1 (5’-TCCTCATGGCATCCTTGTCC-3’) primer and the reverse E1020Kflox-recR11 (5’-TGGTCCACCCGTTGACTCAA-3’) primer by PCR. PCR products were run on a 1% agarose gel. The wild-type allele resulted in a 381 bp band and the recombined p110δ^E1020K^ allele resulted in a 436 bp band.

### Behavioural testing

All mouse behavioural tasks, except the Erasmus ladder, Rotarod and Y-maze, were performed in a behavioural box. This consisted of a 130 x 80 x 80 cm wooden box with a door, lined with 6 mm high-pressure laminate and foam, with a 10 mm Perspex^®^ shelf and standardized white and infrared lights. Metal grooves on the Perspex^®^ shelf assured constant positioning of the testing arenas across experiments. All experiments were recorded with a fixed camera (acA 1300-600gm, Basler AG) positioned above the arenas and operated through the open-source software Bonsai (https://bonsai-rx.org). A frame rate of 25 frames per second (fps) was used for all tests, except for the Grooming assay and the Y-maze, where 30 fps were used. After behavioural testing, video recordings were uploaded to the open-source software OptiMouse (93), where each mouse was tracked and measures such as speed and time spent in regions of interest (ROIs) were extracted. Behavioural tasks were performed as previously described and in the following order: 1) Erasmus ladder (Noldus, Wageningen, the Netherlands) (45); 2) Social interaction (52); 3) Grooming (94); 4) Elevated-plus maze (52); 5) Open-field (95); 6) Marble burying (96); 7) water Y-maze (52); 8) Rotarod (97). The order of the assays was the same for all mice. For detailed information on each assay see Supplemental materials.

### Western blot

Brain tissue was lysed and homogenized in RIPA Lysis and Extraction Buffer (Thermo Scientific™), supplemented with Halt™ Protease and Phosphatase Inhibitor Cocktail (Thermo Scientific™). Protein concentration was determined with Pierce™ BCA Protein Assay Kit (Thermo Scientific™). Protein lysates were mixed with 4X Laemmli Sample Buffer, supplemented with 2-mercaptoethanol (Bio-Rad Laboratories B.V.) and incubated at 100°C for 6 min. Eighty μg (brain and spleen tissue of WT and p110δ^E1020K^ mice) or 40 μg (splenocytes of WT and Pik3cd^−/−^ mice) of lysate were loaded onto 4-15% Mini-PROTEAN^®^ TGX™ Precast Protein Gels (Bio-Rad Laboratories B.V.). Transfer was performed onto Immobilon^®^-P PVDF Membranes (Merck KGaA). Membranes were blocked with 5% BSA (Merck KGaA) in TBS (Merck KGaA) for 1 h and subsequently incubated with anti-PI3K p110δ (D1Q7R) Rabbit mAb (1:1.000, #34050, Cell Signaling Technology, B.V.) in 5% BSA-TBS with 0,1% Tween 20 (TBS-T) (Merck KGaA) overnight at 4°C. Membranes were washed three times with TBS-T and incubated with IRDye^®^ 800CW Goat anti-Rabbit IgG (1:10.000, H + L; LI-COR Biosciences - GmbH) in 5%-BSA-TBS-T for 1 h at room temperature. Membranes were washed three times with TBS-T and imaged in an Odyssey^®^ CLx Imaging System. Afterwards, membranes were incubated with GAPDH (D16H11) XP^®^ Rabbit mAb (1:1.000 dilution, #5174, Cell Signaling Technology, B.V.) in 5% BSA-TBS-T overnight at 4°C. Membranes were washed three times with TBS-T and incubated with IRDye^®^ 800CW Goat anti-Rabbit IgG (1:10.000, H + L; LI-COR Biosciences - GmbH) in 5%-BSA-TBS-T for 1 h at room temperature. Membranes were washed three times with TBS-T and imaged in an Odyssey^®^ CLx Imaging System. Western blots were visualized with Image Studio Lite™ software (LI-COR Biosciences - GmbH).

### Linear discriminant analysis

For multivariate analysis, linear discriminant analysis (LDA) was performed to identify the behavioural features that best separate WT and p110δ^E1020K^ genotypes (58,98). All variables were initially considered for class separation (see pre-processing steps below). Variables that consisted of multiple data points, measured over several sessions (e.g. rotarod data, acquired over the course of 5 days), were reduced to a single value variable by calculating the slope across data points, as this can be interpreted as a learning curve of an animal for a given variable. After pre-processing and validation, LDA was performed with a custom written code. The outcome from the LDA was plotted as LD1 vs LD2, with the contribution of the 10 best variables per LD. All code used to perform the pre-processing steps, validation and LDA is available at https://github.com/BaduraLab/LDA_analysis_2 classes.

#### Pre-processing

Before conducting the LDA, data were pre-processed to comply with the normality assumption by calculating z-scores (98). Z-scores were inspected for every variable and compared with a standard normal distribution. Due to its highly skewed distribution, “Y-maze: reverse II” data were excluded from further analysis. Class approximation of a normal distribution was also assessed by visualising and comparing the z-scored data with a standard normal distribution. Additionally, data points that exceeded 3 scaled median absolute deviations from the median (*isoutlier* function in Matlab) of the corresponding class per variable, were considered outliers and excluded from further analysis. Excluded outliers were interpolated with the mean of their corresponding class per variable (mean interpolation) (98,99).

Next, a correlation matrix with all tested behaviours was generated (Supplementary Fig. 7a) to exclude strongly correlated variables. Inspection of the matrix identified *speed* related variables as strongly correlated (r ≥ 0.86) with measures of *total distance* of the corresponding experiment. Therefore, speed variables were excluded in this step, while distance variables were kept for further analysis, which resulted in 31 behavioural measures included in the LDA. Finally, we applied the Moore-Pseudo Inverse method to allow inclusion of all variables in the analysis by approximating the inverse of the within variance matrix (100). This last step was necessary because one of LDA’s criteria is that the total number of variables analysed must be lower or equal to the total number of samples minus the number of classes (98).

#### LDA validation

To validate the results of the LDA, data was shuffled 200.000 by randomizing data labelling. This number was chosen by shuffling a random dataset *N* times until an error margin of under 5% was achieved, based on the concept of a Monte Carlo simulation (101). With each shuffle, the individual data points were randomly assigned to two equally sized classes. While dominant features can still appear dominant while shuffling, provided that these variables are actually not dependent, exactly equal combinations of contributions were predicted to be low and therefore different from the final LDA results. After shuffling, the first LD1 variable, *Time on ladder (EL)*, appeared 0.21% of the times in 1^st^ place, the second variable, *Light/air ratio (EL)*, was above 18.73% of the times in 2^nd^ place, and the third variable, *Total time (G)*, appeared 1.81% of the times in 3^rd^ place (Supplementary Fig. 7b). The rank sum of the first two features appeared 0.013% of times and the rank sum of the first three appeared 0% of times.

### Statistics

For the analysis of patient visuomotor data, a customized MATLAB script (Mathworks, Natick, MA, USA) was used to visually inspect and analyze all the measured trials. Three outcome measures were considered: 1) Performance - percentage of correctly performed trials; 2) Eye Latency (EL) - time between the presentation of a peripheral stimulus and initiation of the primary saccadic eye movement; 3) Hand Latency (HL) - time between the presentation of a peripheral stimulus and the release of the index finger from the keyboard. The control groups, C1 and C2, were age-matched to patients P1 and P2, respectively. The age and number of control patients is presented in Table 3.

For mouse behavioural data, statistical analysis involving hypothesis testing and group comparison was performed with the Graphpad Prism 8 software. Data sets were first tested for the presence of significant outliers using the Grubbs test, and then for the assumption of normality, using the Shapiro-Wilk test and Q-Q plots. When normality was followed, WT and p110δ^E1020K^ groups were compared with a two-tailed t-test or a 2-way repeated measures ANOVA, depending on the parameters analysed. When data violated the assumption of normality, a two-tailed Mann-Whitney test was performed instead. A mixed effects model was used in place of repeated measures ANOVA when data points were missing or excluded (outliers). The statistical significance threshold was set at p≤0.05. For the analysis of automatically tracked behaviour, body position values were used, except for the “near cup” parameters of the social interaction task. In this case, the nose position was extracted to more accurately represent the interaction between test and novel mice (*sniffing* the novel mouse).

### Study approval

Patient P1 had previously been recruited for a longitudinal, multi-center, cohort study on the causes and clinical manifestations of PID. For this study, approval of the Medical Ethics Committee of the Erasmus University Medical Center Rotterdam had been obtained (MEC 2013-026). Written informed consent was obtained from patients P1 and P2 according to the Declaration of Helsinki.

All experimental animal procedures were approved a priori by an independent animal ethical committee (DEC-Consult, Soest, The Netherlands), as required by Dutch law and conform to the relevant institutional regulations of the Erasmus MC and Dutch legislation on animal experimentation.

## Author contributions

IS, AB, VASHD and JJMP designed and supervised the study. AB, KO, VASHD, JJMP provided resources and acquired funding. VASHD, ORM, SMA and NJMB identified patients and performed clinical diagnosis. IIF, JJMP and AB performed the visuomotor experiments. IS and LW performed mouse experiments. FMPK and HI performed in vitro experiments. IS, IIF, LW and CVDZ performed data analysis. IS performed statistical analysis and prepared figures. IS, AB, VASHD and JJMP wrote the first draft. All authors edited the manuscript. IS and ORM share first authorship. ORM and VASHD provided the clinical characterisation of P1 and P2, whereas IS performed all mouse experiments and coordinated visuomotor data collection and analysis. Because IS drafted the paper, they are listed first.

## Acknowledgements

We thank Peter Katsikis for providing splenocytes of WT and *Pik3cd*^−/−^ animals; Roxanne ter Haar and Elize Haasdijk for the biotechnical assistance; and Chris de Zeeuw for helpful comments and histological reagents. This work has been funded by the Stichting Sophia Kinderziekenhuis Fonds (grant no. S15-07 Genes and Immunity in SCID-F.M.P.K.) and the Dutch Research Council (NWO, ZonMw) Talent Programme Vidi (A.B.)

