## Supplemental materials for "Activated PI3Kδ syndrome, an immunodeficiency disorder, leads to sensorimotor deficits recapitulated in a murine model"

### Supplemental figures

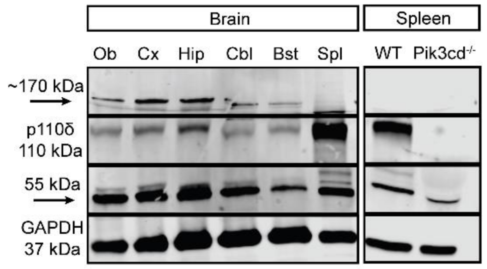


Supplementary figure 1: **p110δ expression in brain tissue**. Representative blot with the expression of p110δ (110 kDa band) across different brain areas in WT mice. Spleen tissue from WT and p110δ$^{E1020K}$ mice was used as a positive control (Spl) while spleen tissue from a Pik3cd$^{-/-}$ mouse was used as a negative control. GAPDH was used as a loading control. Notice the presence of an antibody unspecific band of 55 kDa, and of brain tissue-specific unspecific bands of approximately 170 kDa. Each section presented ("Brain" and "Spleen") represents one individual gel. Ob, olfactory bulbs, Cx, cortex, Hip, hippocampi, Cbl, cerebellum, Bst, brainstem, Spl, spleen. Data from one WT mouse is shown.

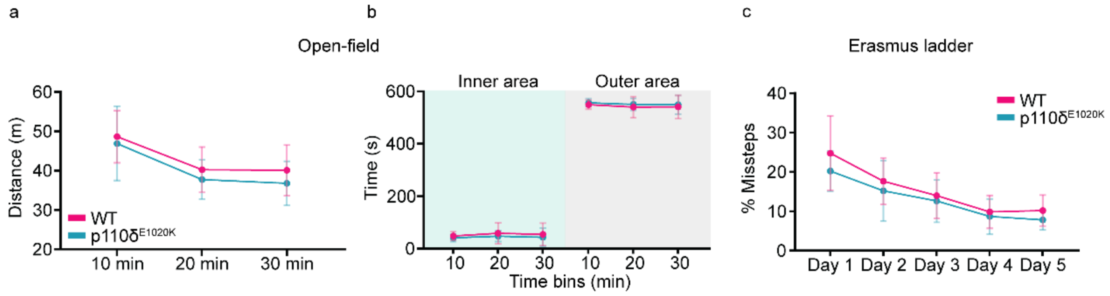


Supplementary figure 2: **p110δ$^{E1020K}$ mice do not show changes in gross motor nor anxiety-like behaviours**. a) Total distance travelled during OF (mixed effects model), and the total time spent by each mouse in the inner (green) and outer (grey) areas of the OF arena (b) were quantified and binned in 10 minute-periods. c) Average percentage of missteps across the 42 daily trials of the Erasmus ladder (mixed effects model). Data are presented as mean ± SD. N = 15 mice per genotype.

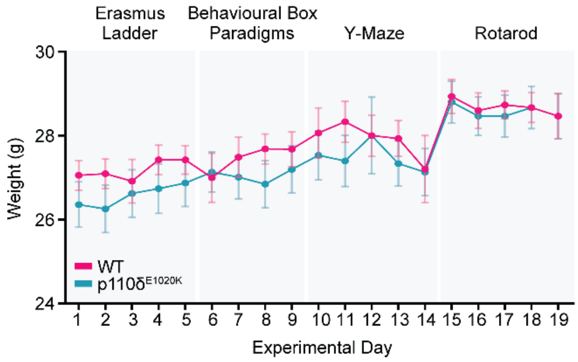


Supplementary figure 3: **WT and p110δ$^{E1020K}$ mice exhibit similar weight progression**. The average weight, in grams, is presented for both genotypes and for each experimental day. Notice the decrement in weight during the Y-Maze week, likely due to increased physical activity (swimming in the maze). Data is presented as mean ± SEM (mixed effects model). n = 15 mice per genotype, except for day 12 (n = 13 WT and 10 p110δ$^{E1020K}$.

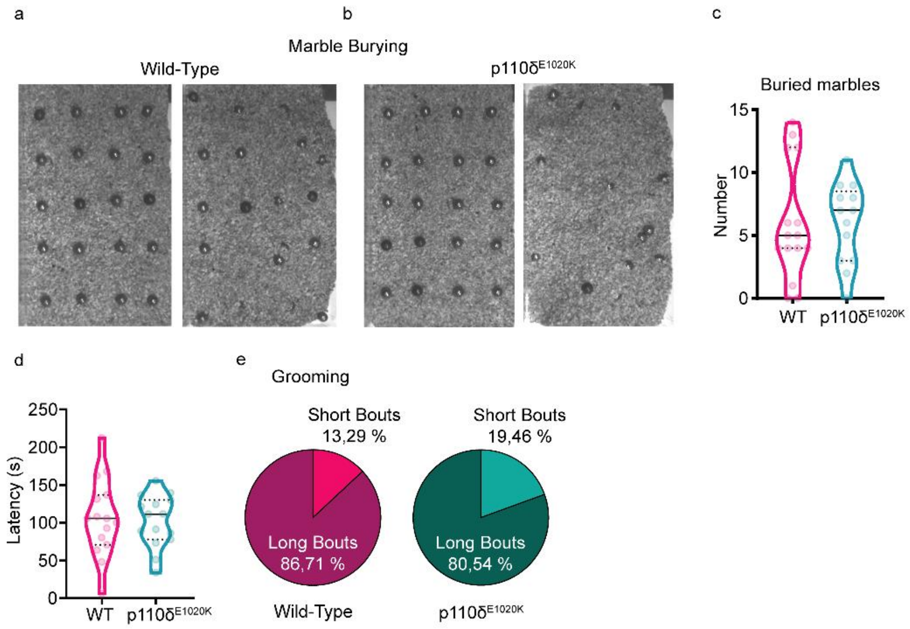


Supplementary Figure 4: **Repetitive behaviour analysis in WT and p110δ$^{E1020K}$ mice**. a-b) Representative pictures of the 4x5 marble set before (left) and after (right) the 30-min marble burying task. c) Quantification of the total number of buried marbles per mouse, using a 50% buried surface cut-off (2-tailed t-test; data presented as median and quartiles, n = 15 WT and n = 13 p110δ$^{E1020K}$ mice). d) Latency, in seconds, to initiate the first recorded grooming bout (2-tailed t-test; data presented as median and quartiles). e) Average percentage of short (<1 second) and long (>1 second) grooming bouts for WT (left) and p110δ$^{E1020K}$ (right) mice. n = 15 mice per genotype, except for c) (see above).

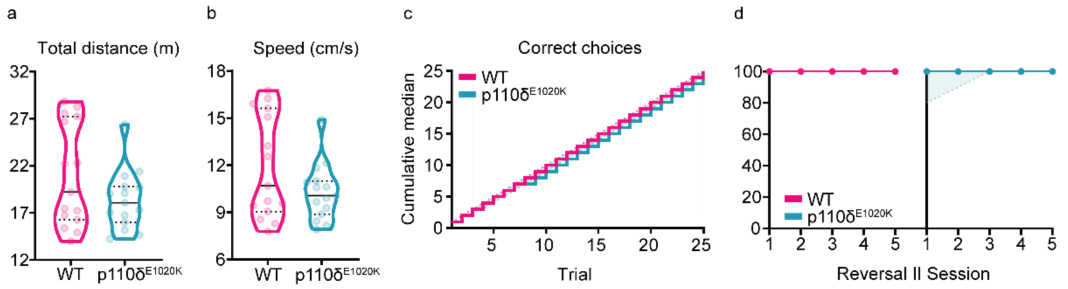


Supplementary Figure 5: **Swimming performance is intact in p110δ$^{E1020K}$ mice**. a-b) Total distance swam (a) and average speed (b) of individual mice during the habituation phase of the Y-Maze. c) Step function with the cumulative median of correct arm choices for the reversal II phase. d) Percentage of correct arm choices for each genotype over the five days of reversal II. Data analysed with 2-tailed Mann-Whitney; n = 15 mice per genotype, except for reversal phases where n = 13.

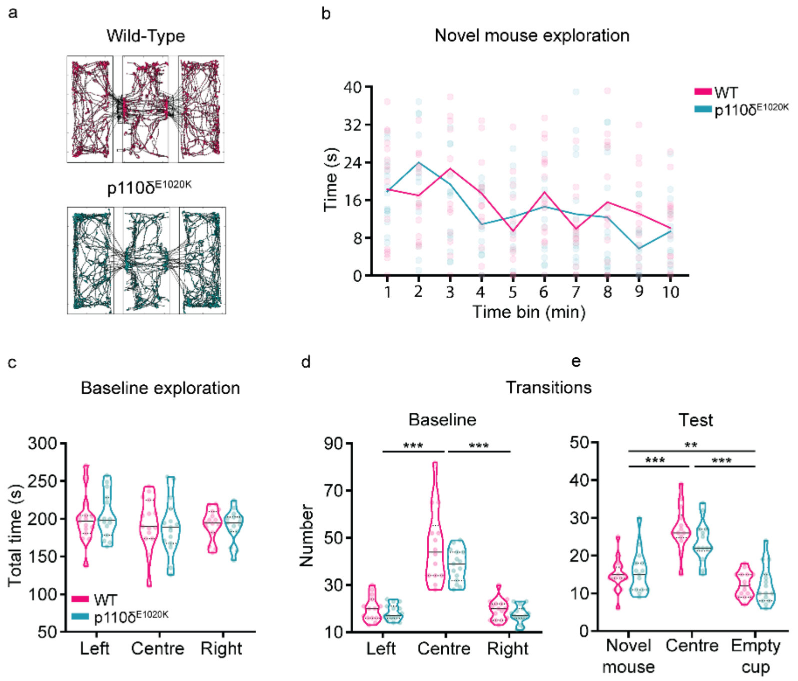


Supplementary Figure 6: **Social behaviour investigation in p110δ$^{E1020K}$ mice**. a) Example of automatically tracked body positions during baseline. b) Total time spent exploring the novel mouse during the 10-minute test phase, split in 1-minute bins (dots represent individual mice and lines connect the mean of each genotype; mixed effects model). c) Total time individual mice spent in each chamber of the SI apparatus during baseline (2-way repeated-measures ANOVA; data presented as median with interquartile range). d-e) Total number of transitions to each chamber made by individual mice during baseline (d) and test (e) (mixed effects model; data presented as median with interquartile range). ** $p \leq 0.01$, *** $p \leq 0.001$. n = 15 mice per genotype.

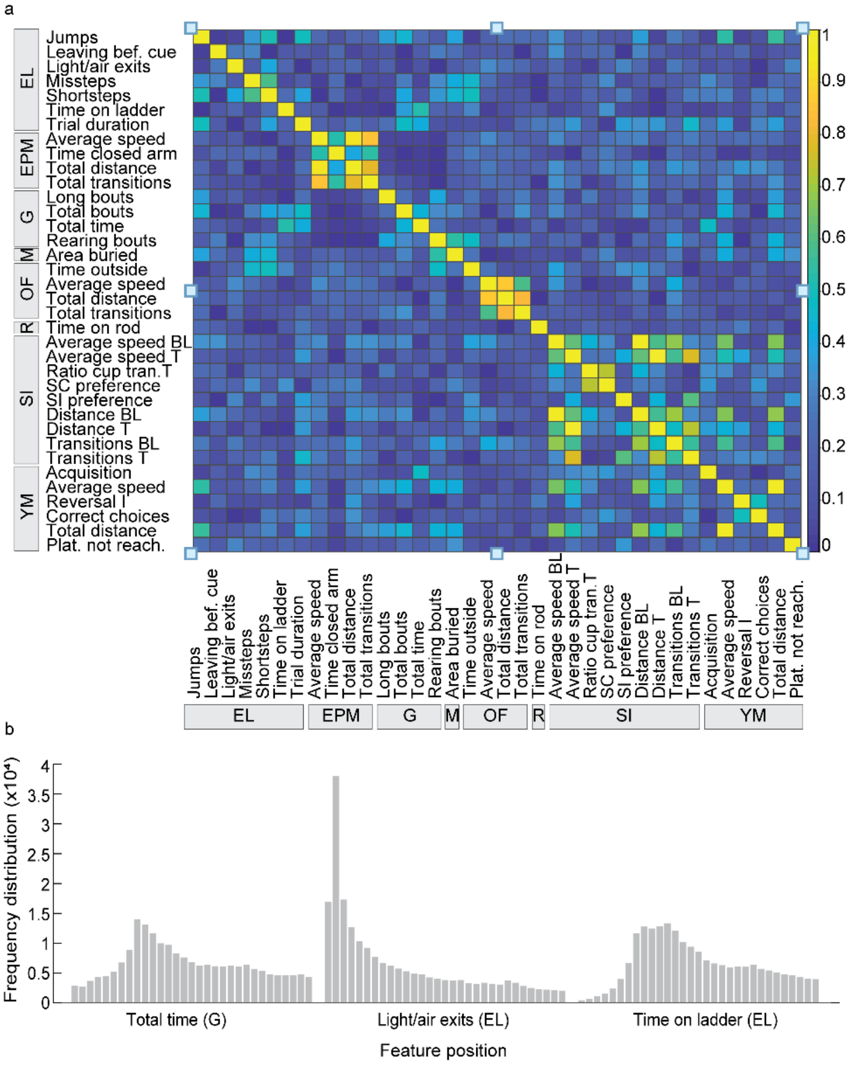


Supplementary Figure 7: **LDA correlation matrix and shuffle validation**. a) After normal distribution validation, a Pearson's correlation matrix with all behavioural variables was plotted to identify strongly correlated variables. b) Example of a shuffling trace with the position distribution of the 3 variables that most contribute to LD1. Data was shuffled 200.000 times to achieve an error margin under the 5%, based on the Monte Carlo Simulation concept. Data represents measurements from 15 WT and 15 p110δ$^{E1020K}$ mice. EL, Erasmus ladder, EPM, elevated-plus maze, G, grooming, M, marble burying, OF, open-field, R, rotarod, SI, social interaction, YM, water y-maze.

**Supplemental methods**

**Behavioural tasks**

*Erasmus ladder*

The Erasmus ladder (Noldus, Wageningen, the Netherlands) was used to assess motor performance and learning (1), over the course of 5 days, with 42 trials per day. The apparatus consists of a horizontal ladder with 37 high and low rungs between two goal boxes. Each trial starts with the mouse inside a dark goal box and a 9-11s waiting period (Fig. 5a). Three seconds after a light is presented, an air puff encourages the mouse to leave the first box and cross the ladder towards the second box. Once the mouse reaches the second box, a new trial is initiated. After 42 trials, the mouse is returned to its home cage and both ladder and boxes cleaned with 70% ethanol. The frequency of before cue, light cue and air cue exits were automatically recorded by the Erasmus ladder software, together with the percentage of backsteps (steps between the current and previous rung), shorsteps (steps between two adjacent rungs), longsteps (steps between the current and second next rung) and jumps (steps between the current and third or above next rung). Absolute totals per day were calculated off line. The percentage of missteps was also calculated and defined as stepping from, to or in a lower rung (1). For one WT and one p110$\delta^{E1020K}$ mouse, data from the second day of the Erasmus ladder is missing due to a power failure during acquisition.

*Social interaction*

Social interaction was evaluated with the three-chamber apparatus as previously described (2). This consisted of a 63 x 42.5 x 21 cm transparent acrylic arena, equally divided in three chambers, separated by two black-opaque movable partitions. On the day prior to testing, age-matched novel mice, of the same strain but different litter, were habituated to wire cups (8 cm diameter, 9.5 cm height) for two periods of 10 min each. On the following day, a test mouse was first allowed to explore the central chamber of the apparatus for 10 min. After this period, the partitions to the right and left chambers were opened, allowing the mouse to explore the full apparatus (*baseline*) during 10 min. The test mouse was then guided to and kept in the central chamber, and an empty wire cup and a wire cup with a novel mouse were placed in a pseudo-randomized fashion in the right and left chambers. The partitions were once again opened and the test mouse was allowed to explore the full apparatus for another 10 min (*test*). At the end of each test and between animals, chambers, partitions and cups were cleaned with 70% ethanol.

*Grooming*

The grooming apparatus consisted of a 30 x 30 x 39 cm white-opaque Perspex® arena. Mice were placed in the arena and allowed to freely explore for 30 min, under dark conditions. At the end of the trial, the arena was cleaned with 70% ethanol and dried before the next mouse was tested. The time and number of grooming events (defined as in (3) was logged in each recorded video using the open-source software BORIS (4).

*Elevated-plus maze*

To investigate the presence of anxiety-like behaviour, an elevated-plus maze (EPM) with two open and two closed arms was used. Each arm was supported by a 30 x 3 cm cylindrical pole and had a 29.5 x 8.5 cm white-opaque acrylic base. Closed arms were additionally surrounded by a 20 cm high black-opaque acrylic wall. At the beginning of each experiment, the test mouse was placed in the centre of the maze and, after 10 min of exploration, returned to its homecage and the maze cleaned with 70% ethanol. The time spent in the closed, open and central areas of the maze, together with the number of entries into each area were calculated.

*Open-field*

To evaluate spontaneous locomotor activity and speed, mice were tracked under light conditions during the open-field (OF) test, in a 50 x 50 x 35 cm white-opaque arena. A test mouse was placed in the centre of the arena and allowed to freely explore for 30 min. After test completion, the mouse was returned to its homecage and the arena cleaned with 70% ethanol. Given that anxiogenic behaviour is associated with increased time spent closer to the walls of the OF (5), ROIs were defined during video analysis (Fig. 4a) to further determine the time each mouse spent in the inner, outer and corner areas of the OF.

*Marble Burying*

The marble burying (MB) test was used to assess repetitive behaviour and anxiety under light conditions (6). A standard 26.6 x 42.5 x 18.5 cm cage (Eurostand 1291H-Type III H) was filled with 4 cm of bedding (Lignocel® Hygienic Animal Bedding, JRS) and 20 blue glass marbles, in 5 rows of 4 marbles, were set on its surface (Supplementary Fig. 4a,b). A test mouse was carefully placed inside the cage and removed after 30 min of exploration. For each mouse, new bedding was used and marbles were cleaned with 70% ethanol.

For the analysis of the surface and number of buried marbles, the open-source programme Fiji was used (7). The first (before testing) and last (after testing) frames of each acquired video were loaded into Fiji and, after scale adjustment, the *freehand selection* tool was used to manually define a ROI around the visible area of each marble. The difference between the visible area on the first and last frames was used to calculate the buried area for each marble and the sum of differences used to calculate the total buried area for each mouse. The number of buried marbles was also determined and a marble was considered buried if its visible surface was reduced in the last frame by 50% or more (6).

*Y-maze*

Flexible learning was tested over the course of 5 days using the water Y-maze, a 3-armed Y-shaped apparatus of white-opaque acrylic (Fig. 5g). Each arm (20 x 32 x 9 cm) has two lateral indentations, 5.5 cm from the centre, that fit a 19.5 x 8 x 0.5 cm white-opaque acrylic wall used during *forced sessions* (described below). The Y-maze was placed in a dark chamber with fixed black poster-board screens on three sides and a movable black curtain on the fourth side. Above the Y-maze, a fixed camera (Sony PS3 Eye) was used to record all trials and enable tracking of distance swam, speed and body position.

In the beginning of each experimental day, the Y-maze was filled with room-temperature water and white paint (Basic color, 21 white 30081, Creall®), to reduce platform visibility. On day 1 (*Habituation*), each mouse swam freely, without a platform, for 3 consecutive 60 s trials, starting from the bottom, then the left and finally the right arm. On day 2 (*Acquisition*), a white-opaque acrylic platform was pseudo-randomly placed in the extremity of the right or left arm. Always starting from the bottom arm, each mouse was placed in the water and allowed to search for the platform over 4 sessions, each with 5 consecutive trials of 40 s. On day 3 (*Test*), the hidden platform was kept on the same side as in day 2. Each mouse performed one single session of 5 consecutive trials, with 40 s per trial. On day 4 (*Reversal I*), the position of the platform was changed to the opposite side (right or left) of the one defined during *Acquisition* and *Test*. Mice searched for the new location of the platform during 5 sessions, each with 5 consecutive trials of 40 s. During session 5, the incorrect arm was blocked (*forced session*), forcing the mouse to eventually swim into the correct arm. On day 5 (*Reversal II*), the protocol used on day 4 was repeated.

Offline manual scoring of correct and incorrect choices was performed for each trial of days 2 to 5. A correct choice was considered if the mouse reached the hidden platform upon the first turn from the bottom arm into the correct (right or left) arm. Data from mice that did not achieve an 80% correct choice rate during *Test*, was not used for analysis of *Reversal I* and *Reversal II* data, as it was considered that these mice did not effectively learn the location of the platform. Two WT and two p110δ$^{E1020K}$ mice met this exclusion criteria.

After each trial, mice were dried with tissue paper and, after each session, returned to their homecage and placed under a heating lamp. At the end of each day, the Y-maze was emptied, cleaned with tap water and 70% ethanol, and dried.

*Rotarod*

Motor performance was assessed with the rotarod (8). We used a five-day accelerating rod protocol, each with 4 non-consecutive trials per mouse per day. From days 1 to 4, the speed of the rotating rod was accelerated to 40 rpm while, on day 5, maximal speed was increased to 80 rpm. The time each mouse stayed on the rotarod was counted by the apparatus' stopwatch and manually recorded for each trial. Between trials, mice were given a 1 h resting period, and the rod and separating walls were cleaned with 70% ethanol and dried with paper towels. A trial was considered finished when the mouse fell off the rod, grabbed the moving rod performing a 360º turn without actually walking on the rod or reached a maximum time of 300 s on the rod. The average time per trial across the five days and the total time spent on the rod were calculated.
